# Sortase A-based post-translational modifications on encapsulin nanocompartments

**DOI:** 10.1101/2024.03.05.583518

**Authors:** Seyed Hossein Helalat, Yi Sun, Rodrigo Tellez

## Abstract

Protein-based encapsulin nanocompartments, known for their well-defined structures and versatile functionalities, present promising opportunities in the biotechnology and nanomedicine area. In this investigation, we effectively developed a sortase A-mediated protein ligation system in *E. coli* to site-specifically attach target proteins to encapsulin, both internally and on its surfaces without any further *in vitro* steps. We explored the potential applications of fusing sortase enzyme and a protease for post-translational ligation of encapsulin to a green fluorescent protein (GFP) and anti-CD3 scFv. Our results demonstrated that this system could attach other proteins to the nanoparticles’ exterior surfaces without adversely affecting their folding and assembly processes. Additionally, this system enabled the attachment of proteins inside encapsulins which varied shapes and sizes of the nanoparticles due to cargo overload. This research opens up avenues for further exploration of the applications and engineering possibilities of encapsulins in the rapidly advancing field of nanotechnology.

## Introduction

Nanoparticles have revolutionized biomedical research and nanomedicine by offering versatile surfaces and functional cores with unique chemical and physical properties. Diverse nanomaterials like lipids, quantum dots, polymers, and metal oxides have been harnessed for their drug-delivery capabilities and therapeutic potential.^1^

Among them, encapsulin nanocompartments, as protein-based nanoparticles, represent a class of supramolecular structures with spherical and hollow architectures, exhibiting highly symmetric and uniform sizes.^2^ Encapsulins, widespread in bacteria and archaea, are homooligomers, typically composed of a single subunit, assembled into large 60-mer, 180-mer, or 240-mer assemblies, that encapsulate cargo proteins and play a crucial role in segregating enzymes, ions storage, and preventing toxic side reactions.^3^

Encapsulin nanocompartments offer versatile applications ranging from vaccine development to targeted drug delivery.^4–6^ Encapsulins can act as efficient nanocarriers,^7^ and their inherent properties make them promising candidates in the field of nanomedicine, highlighting the potential for enhanced therapies and diagnostics.^8^

In this context, *Quasibacillus thermotolerans*’ large encapsulin compartment has garnered attention. This 240-mer encapsulin system is the largest identified nanocompartment and, self-assembles into a thermostable protein compartment, which in *Q. thermotolerans* plays a role in storing an exceptionally large quantity of iron. ^9,10^

The manipulation of protein-based nanoparticles is crucial for obtaining nanoparticles tailored for various uses. It is essential to have control over both the interior and exterior features. Various studies have been conducted to alter these particles, involving adjustments to pore sizes, charges in different regions, and modifications to the exterior side. These modifications aim to streamline the purification of nanoparticles and enable the attachment of functional molecules onto their surfaces, like targeting ligands, antigens for nano vaccines,^5^ and other proteins for bioimaging or biosensor production aims.^2,11,12^ However, direct genetic fusion or chemical conjugation of whole proteins or domain ligands to nanoparticle’s protein backbone is challenging due to potential improper protein folding and aggregation. ^5,13,14^

In this regard, chemical linkers have certain drawbacks, including imprecision, variability, potential steric hindrance, complex synthesis, and high cost.^15,16^ Similarly, biological methods such as the SpyTag-SpyCatcher system are not without limitations. A significant drawback is that modifications required at the protein termini may potentially alter the protein’s structure or function, particularly given the size of SpyCatcher (approximately 13 kDa). The efficiency and specificity of binding can vary depending on the proteins involved and reaction conditions, presenting challenges in achieving high efficiency and specificity.^5,17,18^ Additionally, the use of this system may increase the immunogenic effect of nanoparticles.^19^

In recent years, enzymatic conjugation systems like Sortase A (SrtA) have emerged as a robust approach for efficient protein conjugation and post-translational modifications.^15^ SrtA catalyzes a transpeptidation reaction that covalently links a peptide motif (LPXTG) with a poly-glycine motif (GG) on the target protein. SrtA-mediated ligation does not require adding large domains at the protein termini, minimizing the risk of altering protein function and it offers high specificity, resulting in site-directed conjugation, which is crucial for maintaining the protein’s functionality.^20^ Additionally, SrtA-mediated conjugation occurs under normal cell physiological conditions, preserving the native structure and activity of the proteins involved, making it a suitable option for complex protein-protein conjugations. ^21,22^ These advantages make SrtA an attractive tool for conjugating proteins and post-translational modifications of encapsulin nanocompartments to attach target proteins on the surface of the nanoparticles after their translation and folding.

Additionally, to uptake recombinant proteins into encapsulin nanoparticles, it is feasible to introduce a specific cargo-loading peptide to the recombinant protein. ^23^ However, up-to-date, enzymatic (Sortase-based) approaches for engineering the interior side of these particles and attaching different proteins have not been explored.

In this investigation, we explored the use of the SrtA enzyme for post-translational modifications on both the interior and exterior surfaces of *Q. thermotolerans* encapsulin nanoparticles in *E. coli*. To achieve this, we generated a fusion protein consisting of srtA and Tev protease (to create a poly-glycine motif), facilitating the ligation of two proteins. Subsequently, we examined the system’s efficacy in attaching GFP both inside and on the surface of encapsulin nanoparticles. Our findings indicate that SrtA offers a versatile and efficient method for protein conjugation, eliminating the need for additional *in vitro* steps.

## Materials and Methods

### Plasmid design and preparation of genes

To establish the post-translational modifications, we synthesized srtA with 4 mutations, tev, and *Q. thermotolerans* encapsulin genes. These genes were then cloned into the pETDuet™-1 vector using the Gibson assembly kit. Afterwards, the tev protein was fused to srtA by a flexible linker (2x GGGGS) to make a fusion assembly protein (SrTev). Then, Ribosomal Binding Sites (RBS) sequences were replaced to decrease the expression of the assembly genes.

As cargo proteins, super fold green fluorescent protein for the interior and exterior attachment, and anti-CD3 single-chain variable fragment (scFv) for exterior attachment were used.

For conjugation using srtA and tev, a specific LPETG peptide sequence was introduced at the C-terminus of the target proteins during recombinant protein expression, Enc-srt and GFP-srt. Also, a specific tev digestion peptide sequence (<u>ENLYFQG</u>GGGS) was introduced in the N-terminus of the proteins to generate the poly-glycine motif in the N-terminus of encapsulin (Enc-GG), GFP (GFP-GG), and anti-CD3 scFv (scFv-GG) . The term GG was used to describe the poly-glycine motif which represents the four glycine amino acids downstream of the cleavage site.

In addition, to make a control for GFP uptake by encapsulin nanoparticles, a native *Q. thermotolerans* encapsulin was cloned into the expression vector plus a GFP gene fused to the specific Cargo-Loading-Peptide (CLP) sequence (Table S1). Moreover, we added the scFv-GG gene into this construct to generate fluorescent nanoparticles with loaded GFP-CLP proteins and anti-CD3 scFv on the surface for targeting the Jurkat cells. Notably, a longer flexible linker (4x GGGGS) was added to the N-terminus of the scFv proteins for their attachment on the surface, which allows for flexible movement.

### Gene Expression and purification of proteins

To facilitate plasmid propagation, we employed *E. coli* DH5α strain, which was cultured in LB medium (10 g/L tryptone, 5 g/L NaCl, and 5 g/L yeast extract) at 37°C with continuous shaking at 250 rpm. The resulting constructs were verified through sequencing, followed by transformation into the *E. coli* BL21 (DE3) strain for gene expression.

For protein expression, 2 mL of the overnight culture was transferred to 200 mL of fresh media containing 100 µg/mL ampicillin and incubated at 37°C (250 rpm) for 3 h. Once the OD600 reached approximately 0.8, IPTG was added to a final concentration of 0.5 mM to induce protein expression. Then, to evaluate the assembly proteins (SrtA and Tev) activity, the cells were incubated overnight at 18, 25, 30, and 37°C. For protein purification, pellets from the 200 mL cultures were obtained, re-suspended in PBS, and lysed via sonication. Lysate was centrifuged and supernatant was used for protein purification employing 500 µL of HisPur™ Ni-NTA Resin and Pierce™ Spin Columns (Thermo Fischer Scientific), following the manufacturer’s instructions. Of note, 50 mM and 500 mM imidazole were used as the washing buffer and elution buffer, respectively, according to in-house optimization tests. After purification, to increase nanoparticles’ purity, size exclusion chromatography (SEC) was employed using a HiPrep 16/60 Sephacryl S-500 HR column (GE Healthcare) in PBS buffer (1 mL/min) on an AKTA Pure machine. Then, nanocapsule fractions were concentrated using Amicon Ultra-15 Centrifugal Filter Units with Ultracel-100 membrane (Millipore), then diluted in 2 mL of 20 mM Tris buffer at pH 8. Ion-exchange chromatography using a HiPrep DEAE FF 16/10 column (GE Healthcare) resulted in the fully purified particle sample for further analysis. The gradient used for ion-exchange was 100% A for 0–70 mL, 100% A to 50% A + 50% B for 70–200 mL, 100% B for 200–300 mL, 100% A for 300–400 mL; where A is 20mM Tris pH 8, B is 20mM Tris pH 8 with 1M NaCl (flow rate: 3 mL/min). Subsequently, the buffer of the proteins was exchanged, and the concentrations were adjusted using Amicon® Ultra-15 Centrifugal Filter Units (Merck) with 100 kDa cut-off, employing PBS solution containing 0.5 mM DTT, and 5 mM MgCl2.

### Post-translational modifications assessment

Protein yield, integrity, and protein ligation with GFP were assessed by analyzing 5 µL of the purified proteins using SDS-PAGE. The ligated encapsulin protomers should show larger bands and the efficiency of the ligation was evaluated by measuring the ratio between native and ligated encapsulin protomers using gel densitometry with ImageJ software. Also, the fluorescence of samples with the same concentration (100 µg) was measured by a plate reader (excitation 485 nm, emission 512 nm) to assess the GFP emission from the ligated samples.

Also, for the encapsulin nanoparticles with scFv proteins on their surface, due to the low concentration of the encapsulin+scFv ligated band on the gels, we utilized western blotting. Briefly, the cells were lysed by adding Laemmli buffer containing 10 mM DTT, followed by boiling. Protein extract samples were loaded onto an SDS-PAGE gel and transferred to a polyvinylidene fluoride (PVDF) membrane. The PVDF membrane was prepared by treating it with ethanol and water. The transfer sandwich, including the membrane and gel, was placed in the transfer buffer and transferred at 140 mA for 90 min. The membrane was then blocked with skim milk in PBS-Tween, washed, and incubated with a 6x-His-Tag monoclonal antibody (Invitrogen) diluted 2500-fold in PBS-Tween overnight. After washing, a chemiluminescence reaction was performed using Pierce™ ECL Western Blotting Substrate (Thermo Fisher Scientific). Finally, the membrane was imaged using a CL detector.

### Cell culture and targeting the cells

Jurkat cells were cultured at 37°C with 5% CO__2__ in RPMI medium supplemented with 10% fetal bovine serum (FBS) and 1% PenStrep (100 U/mL penicillin and 100 μg/mL streptomycin. The cell cultures were tested for mycoplasma contamination before the experiments.

To target the Jurkat cells with encapsulin nanoparticles carrying GFP internally and harboring the anti-CD3 scFv on the surface, 20 µg of nanoparticles were added to Jurkat cells and incubated at 37°C with 5% CO__2__ for 2 hours. Subsequently, treated cells were washed 3 times with PBS, and the binding ability was examined by fluorescence microscopy.

### Transmission electron microscopy (TEM)

To visualize the morphology, size, and state of encapsulin assembly, TEM was performed using a Tecnai G2 T20 (200 kV accelerating voltage). Briefly, encapsulin samples (0.2 mg/ mL) were adsorbed onto carbon 200 mesh copper grids for 3 min. Prior to imaging, samples were washed with ultrapure water and then negatively stained for 2 min using VitroEase™ Methylamine Tungstate Negative Stain (Thermo Scientific Co.) and allowed to dry for 15 min. ImageJ software was used to analyze the TEM images.

### Size determination of encapsulin-GFP nanoparticles

Purified encapsulin nanoparticles with GFP proteins ligated either in their outer surface (C-ter) or internally (N-ter), were suspended in PBS and sonicated in a water bath for 30 s. Then, by using a Nanoparticle Tracking Analysis (NTA) the size of nanoparticles was measured. To this end, 1 mL of each sample solution was injected into the NTA instrument, and particle size was determined for 30 s with a flow rate of 100µL/min by triplicate.

## Results and Discussion

### Manipulations of encapsulin protomers

The required genes including encapsulin from *Q. thermotolerans*, GFP, srtA, and tev were successfully synthesized and cloned into the pETDuet™-1 vector. Transformation into *E. coli* BL21(DE3) cells resulted in the successful expression of all required proteins upon IPTG induction. SDS-PAGE analysis confirmed the expression of the desired protein. Also, purification steps were performed through Ni-NTA resins, SEC, and ion exchange chromatography (Figures S1a-b).

The protein structure of the *Q. thermotolerans* encapsulin (PDB 6NJ8) shows the N-terminal of the protomers is located inside of the nanocompartment and their C-terminal is accessible on the surface of the nanoparticles.^9,24^

As our aim was to perform post-translational modifications within *E. coli* rather than *in vitro* (tubes), for srtA-mediated ligation, it was essential for the C-terminal of Enc-srt or GFP-srt (the first protein) to contain the LPETG motif, and the N-terminal of Enc-GG or GFP-GG (the second protein) to have the poly-glycine (GG) motif that contained four glycine amino acids in this study. Given that a methionine is naturally present in the N-terminal of proteins during gene expression in *E. coli*, we introduced a Tev protease recognition site at the N-terminal of Enc-GG and GFP-GG to generate a poly-glycine motif after digestion. Then, this poly-glycine segment could be ligated to the LPETG motif in the first protein (Figure 1). Our results indicated that the presence of the LPETG motif and Tev recognition site, whether in the C-terminal or N-terminal, does not adversely affect the encapsulin protomers and their assembly, and protomers were soluble and could assemble properly. This lack of negative impact could be attributed to the small size of these motifs. Furthermore, the inclusion of these short motifs did not demonstrate adverse effects in the N-terminal and C-terminal of the GFP protein as well.

**Figure 1.**
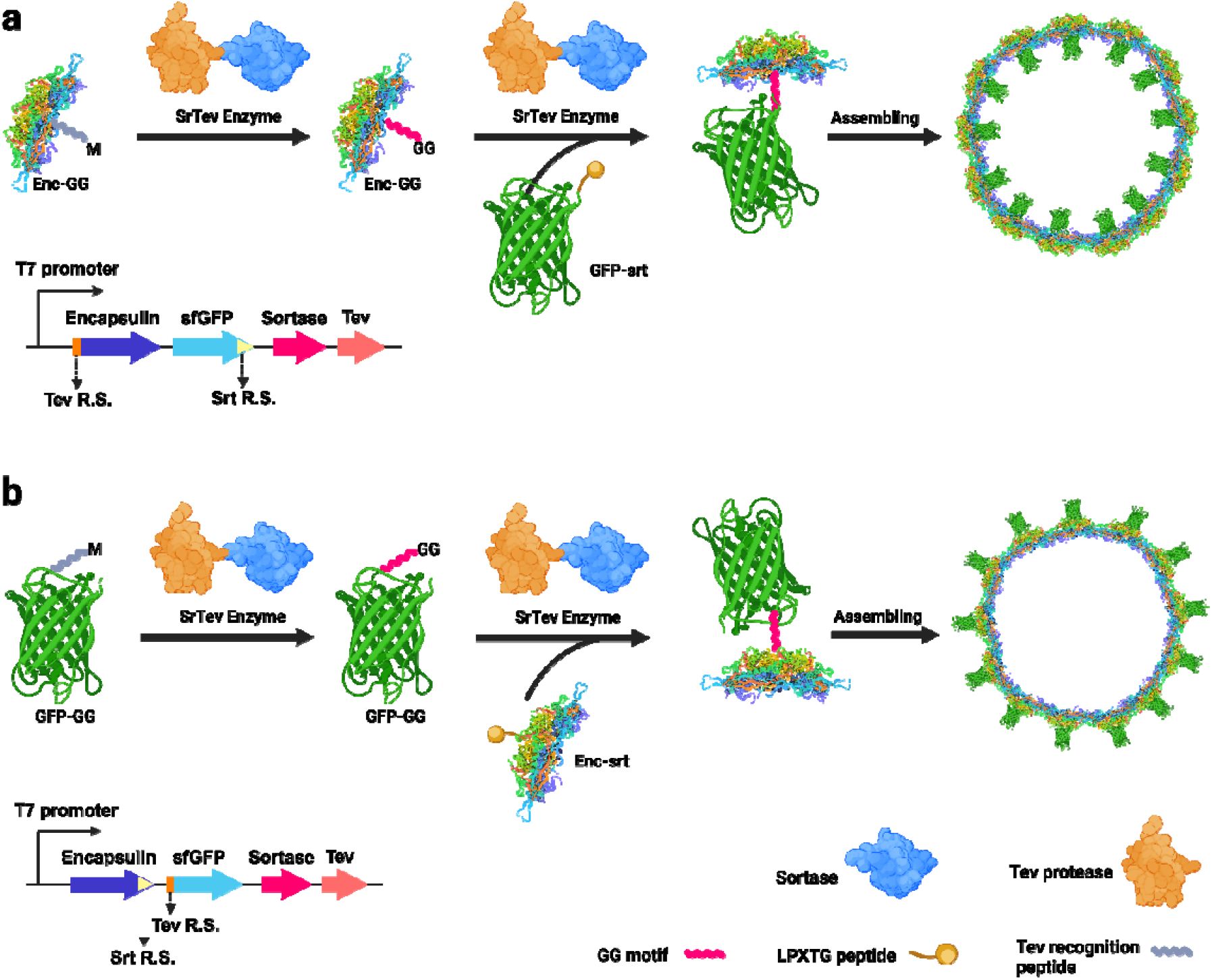
Post-translational modification of *Q. thermotolerans* encapsulin. (a) Incorporating cargo protein inside encapsulin nanoparticles. In this process, the Tev recognition site in the N-terminal of Enc-GG undergoes cleavage by Tev protease, resulting in the generation of the GG motif. Subsequently, GFP-srt is ligated to the newly formed GG motif. (b) Attaching cargo protein to the surface of encapsulin. To achieve this, the Tev protease is employed to generate the GG motif in GFP-GG, which is then attached to the LPETG motif in the C-terminal of Enc-srt by the SrtA enzyme.

### Sortase activity

Efficient srtA-mediated ligation was achieved by genetically fusing target proteins with a peptide containing the LPETG motif recognized by srtA, along with a recognition site for the Tev enzyme to generate a GG site at the N-terminus of the proteins. SDS-PAGE results confirmed the successful ligation of the target protein to encapsulin, and our results demonstrated that approximately half of the encapsulin protomers in purified encapsulin nanocompartments were ligated (Figure 2a).

**Figure 2.**
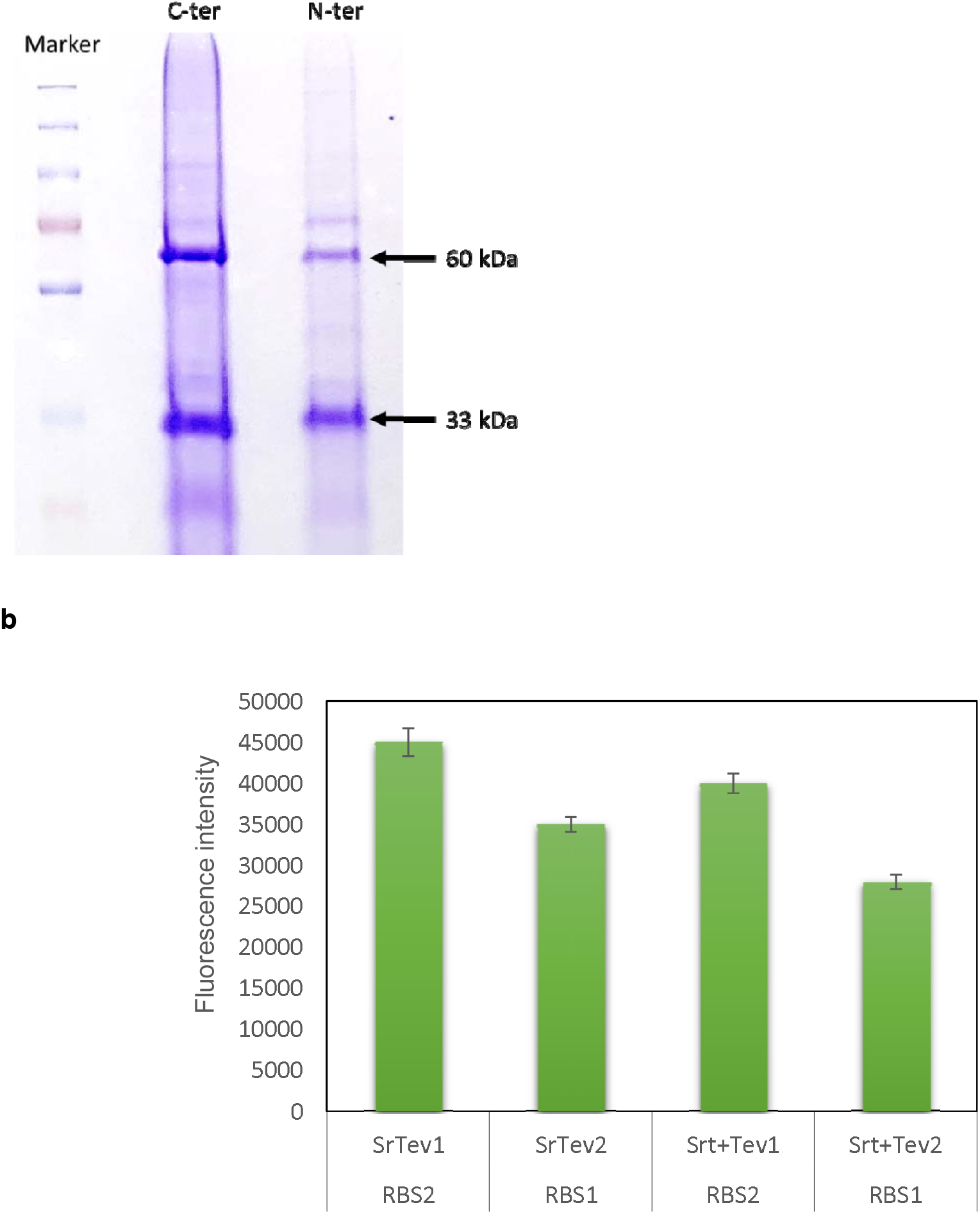
Sortase-based ligation of encapsulin and GFP in *E. coli*. (a) SDS-PAGE image of ligated encapsulin nanocompartments after purification. The image illustrates the attachment of GFP protein (approximately 27 kDa) to the encapsulin protomer (33 kDa), resulting in the observation of a ligated protein with a size of 60 kDa in the gel. From left, well number 2 (C-ter), corresponds to encapsulin nanoparticles with GFP on their surface, while well number 3 (N-ter) represents nanoparticles with GFP inside. (b) The fluorescence intensity was measured for purified nanoparticles at a concentration of 100 μg/mL. SrTev-RBS2 signifies ligation facilitated by the SrTev fusion enzyme with a weak RBS, while SrTev-RBS1 represents ligated nanoparticles through SrTev with a strong RBS (pETDuet™-1 vector RBS). In Srt+Tev samples, individual enzyme forms were employed.

Furthermore, a fusion protein of srtA and Tev exhibited activity in both digesting the Tev recognition motif in the N-terminal of proteins and transpeptidation of the LPETG peptide in the C-terminal of encapsulin and GFP proteins. Our results indicated that the SrTev fusion protein’s efficiency surpassed that of separate enzymes at lower enzyme concentrations. To assess this, we utilized low-translation RBS sequences upstream of fusion and separate genes, comparing their ligation efficacy. The results revealed that the fusion protein exhibited higher performance, potentially attributed to the increased accessibility of digested segments to the srtA enzyme (Figure 2b).

To investigate the *in situ* activity of the proposed system, we tested different temperatures for the modification of *Q. thermotolerans* encapsulin, expressing proteins at 18, 25, 30, and 37 degrees Celsius. Our findings showed consistent activity of the system across all selected temperatures. Moreover, we observed that reducing the expression level of sortase and Tev enzymes compared to encapsulin and GFP, achieved through the use of low-expression RBS sequences, increased conjugation efficiency. This phenomenon may be attributed to adjusting the ligation time ratio to the proper folding and assembly of encapsulin proteins, along with a decrease in the metabolic burden associated with the excess production of srtA and Tev (Figures 2b and S1c).

### Encapsulin exterior side modifications

Previous studies have shown that it is possible to fuse short peptides (5-25 aa) to the N-terminal ^25^ or C-terminal of the encapsulin proteins and the particles could be assembled correctly. Also, fusing short sequences into the E-loop of the encapsulins is possible.^18,26^ However, fusing larger proteins can cause aggregation or lack of assembly ability of encapsulin protomers. Nevertheless, it is possible to attach other molecules and proteins after translation of the encapsulin protomers or on the assembled nanoparticles. ^2,17,19^

In our study, the GFP proteins were attached to the surface (C-terminal) of the encapsulin nanocompartments by the SrTev fusion protein. Based on our findings, in the best-optimized conditions, such as IPTG concentrations, induction temperature, and replacement of RBS sequences, in the purified nanoparticles, 40% of the protomers were ligated to the GFP (Figures 2a and S1c). This phenomenon could be due to the spatial hindrance of the GFP proteins on the surface. Also, our transmission electron microscopy (TEM) findings indicate that utilizing the proposed system to attach GFP to the surface of encapsulin nanocompartments does not impact the shape and backbone size of these nanoparticles (Figures 3a and S2a). However, NTA results for the size of the nanoparticles indicated that the size of the particles was around 46 nm, which was due to adding the GFP proteins all around the nanoparticles (Figure 3a).

**Figure 3.**
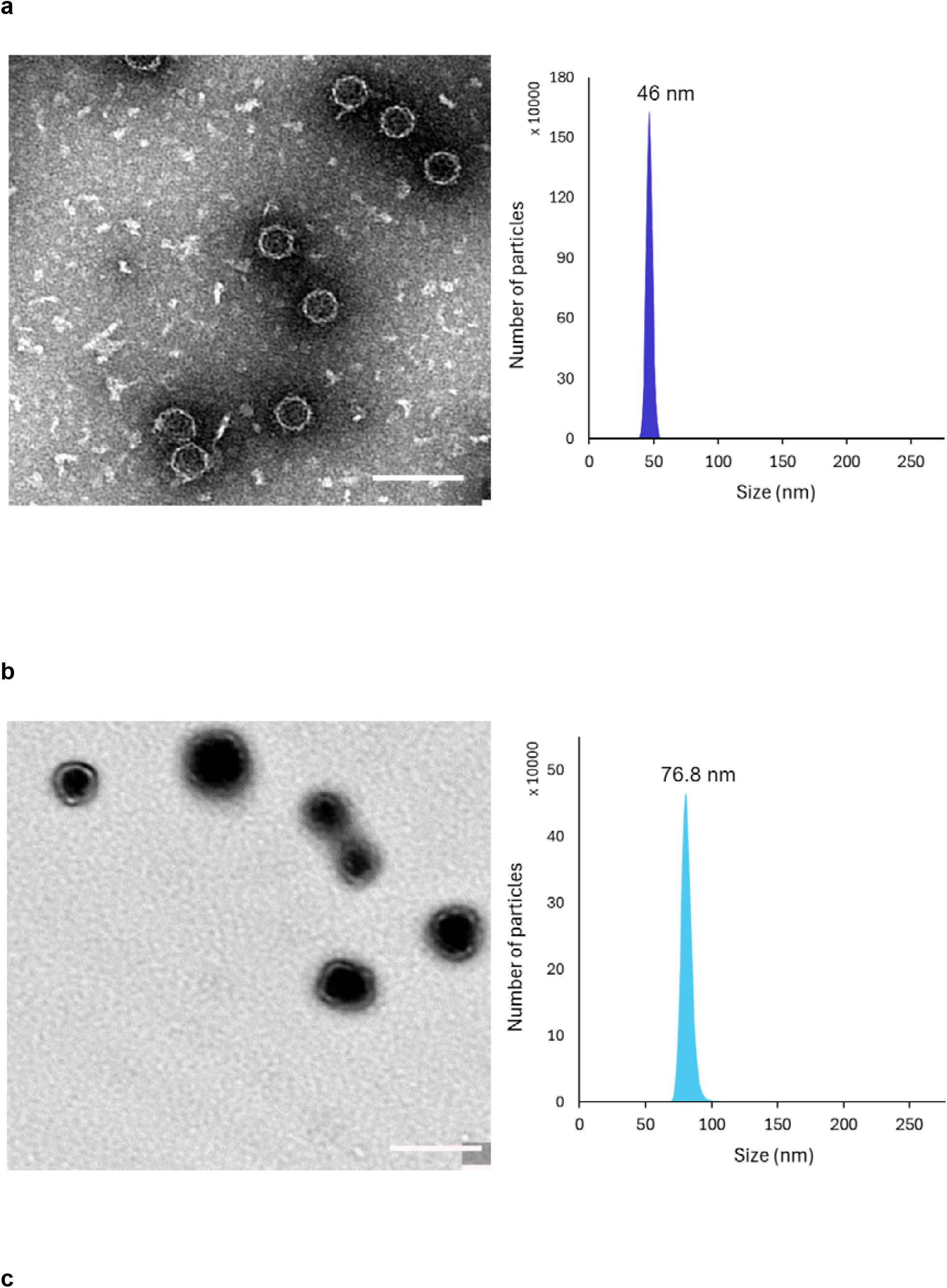
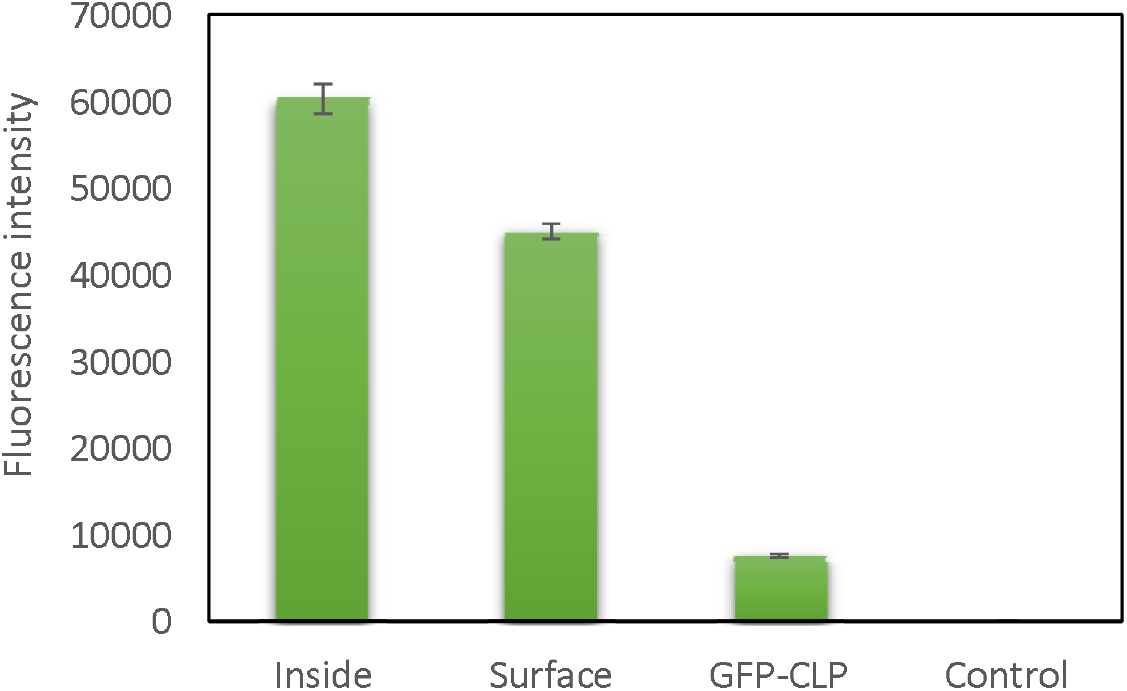
TEM image and NTA analysis results of the post-translational modified encapsulin nanoparticles. (a) Encapsulin nanoparticles with GFP proteins ligated to the surface exhibited consistent shapes with a backbone size of 42 nm (TEM image) and full size of 46 nm (NTA analysis). (b) Encapsulin nanoparticles with GFP ligated internally showed varied shapes and sizes ranging due to cargo overload (TEM image) and an mean size of 76.8 nm in NTA analysis results. The shapes deviate from the typical spherical form, as evident in the image. The scale bar represents 100 nm. (c) Fluorescence intensity of purified nanoparticles at a concentration of 100 µg/mL. From left to right: the first sample represents encapsulin nanoparticles containing GFP internally (Inside), the second shows nanoparticles with GFP ligated on their surface (Surface), and the third indicates encapsulin nanoparticles harboring GFP with fused CLP (GFP-CLP). The control sample consists of purified encapsulin without the presence of the srtA enzyme.

These findings showed the potential of employing the sortase-based ligation system to attach various proteins to encapsulin nanoparticles. This approach mirrors previous successful studies demonstrating the attachment of proteins to other nanoparticle platforms, such as E2 and metal-based nanoparticles. ^27,28^

### Encapsulin interior side modifications

While various techniques, such as chemical linkers and the spytag-spycatcher system, have been explored to attach molecules on the surface of encapsulin nanoparticles, in this investigation, we explored the feasibility of attaching proteins inside the encapsulin nanoparticles by the designed system. The standard method for encapsulating proteins and other molecules into these nanoparticles involves using CLP sequences. These sequences possess a specific affinity to the interior side of encapsulin through hydrophobic and ionic interactions. While numerous studies have examined the assembly of therapeutic proteins and enzymes with CLP motifs, none have investigated the possibility of attaching proteins internally without causing protein aggregations and issues with nanoparticle assembly. ^9,23^

Employing the srtA system for this purpose demonstrated that post-translational attachment of proteins within encapsulins can enhance the quantity of loaded proteins. However, an overload of cargo did impact the shape and size of the nanoparticles.^4^ Our results revealed an eightfold increase in GFP loading using this system compared to the use of CLP (Figure 3c). Nevertheless, varied shapes and sizes of encapsulin nanoparticles were observed probably due to cargo overload. TEM results revealed varying sizes of the nanoparticles, and NTA analysis indicated a mean particle size of approximately 76.8 nm. Additionally, the spherical shape of the particles sometimes appeared elliptical (Figure 3b).

Previous studies on cargo overload with CLP in *Myxococcus xanthus* encapsulin nanoparticles exhibited a similar phenomenon, and Seokmu Kwon and his colleagues demonstrated that shapes could be distorted or form fusion shells, defined as two-lobed particles.^4^

Furthermore, the SDS-PAGE examinations of the purified encapsulin nanoparticles for the presence of trapped SrtA and Tev enzymes did not show any identifiable enzymes on the gel. This suggests the probable absence of trapped enzymes within the nanoparticles. It can be inferred that the ligation of the two proteins happens when the protomers are in their individual, unassembled states, rather than during the process of nanoparticle assembly.

### Expression and attachment of anti-CD3 scFv

To demonstrate the versatility of the system, we expressed an anti-CD3 scFv antibody and conjugated it to the surface of encapsulin nanoparticles by our system for targeting Jurkat cells, serving as a T-cell model. Also, we encapsulated GFP-CLP proteins within the nanoparticles to produce fluorescently labeled particles (Figure 4a).

**Figure 4.**
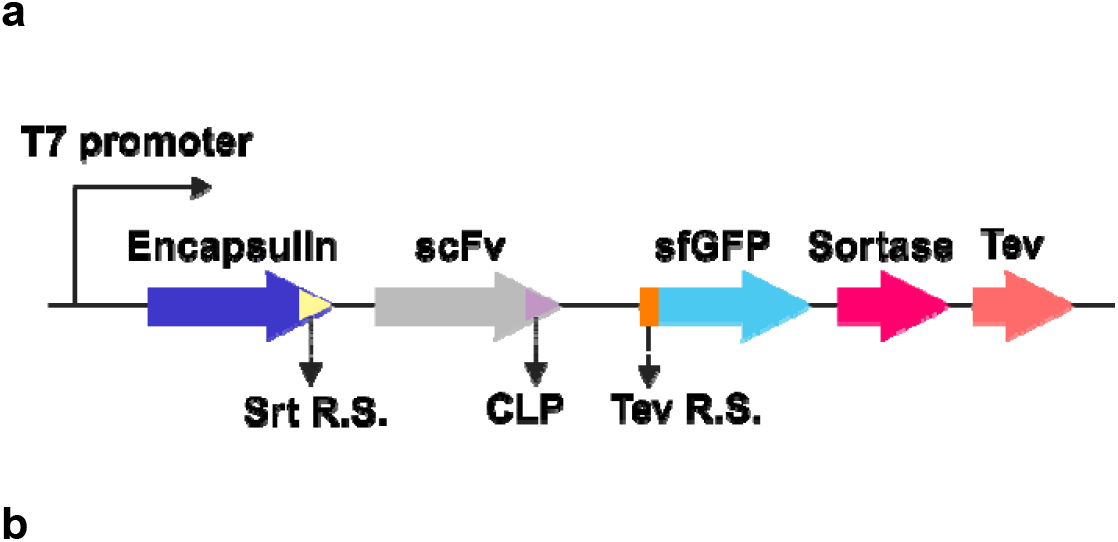
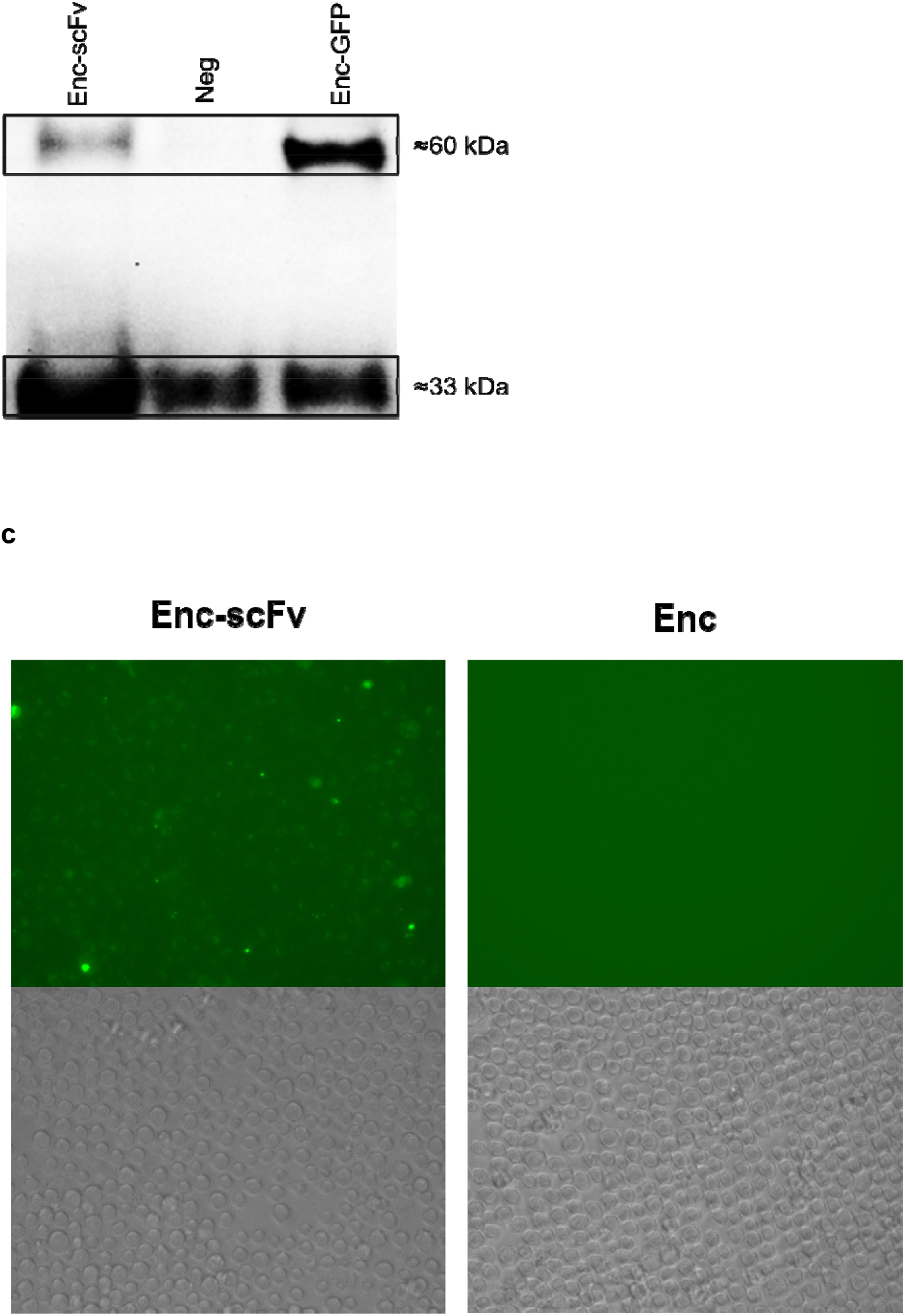
Assessment of the system for attaching anti-CD3 scFv proteins on the surface of the encapsulin nanoparticles. (a) The map of the constructed plasmid for co-expression of the scFv with the developed assembly system. (b) Western blot image for the purified encapsulin nanoparticles with scFv (Enc-scFv) and GFP (Enc-GFP) on their surface. The negative control (Neg) was the normal encapsulin nanoparticles. The variation in size between the ligated proteins in Enc-scFv and Enc-GFP is attributed to the 3.5 kDa difference in the scFv, which is larger than the GFP protein. (c) Fluorescence microscopy images of Jurkat cells treated with encapsulin nanoparticles carrying scFv on their surface (Enc-scFv) and normal encapsulins (Enc). The nanoparticles contained GFP-CLP protein, and green fluorescence indicative of binding could be observed.

Our expression of scFv in *E. coli* showed a very low expression level and the formation of inclusion bodies. The band corresponding to scFv was not detectable on SDS-PAGE gels after purification. Similarly, co-expression of scFv with our assembly system yielded a weak band corresponding to the size of ligated encapsulins with scFv. To validate the attachment of scFv to the surface of the encapsulin nanoparticles, we employed western blot analysis, and its results confirmed the successful ligation (Figure 4b).

To further explore these nanoparticles, we introduced fluorescent encapsulin nanoparticles with scFvs on their surfaces to Jurkat cells. Following a 2-hour incubation, the cells were washed, and subsequent evaluation was performed using fluorescent microscopy. The results demonstrated successful binding of nanoparticles to Jurkat cells, effectively labeling the cells and turning them green upon binding. As a negative control, we added normal encapsulins with GFP-CLP inside, but no cell labeling or fluorescence was observed (Figure 4c).

The obtained results could open a new avenue to nanoparticle and antibody conjugation without the need for further *in vitro* (tube) steps.

## Conclusion

This study successfully employed a SrtA-mediated protein ligation system to achieve site-specific post-translational modifications on both the interior and exterior surfaces of *Q. thermotolerans* encapsulin nanocompartments in *E. coli*. The results demonstrated the versatility and efficiency of the proposed system, eliminating the need for additional *in vitro* (in tubes) steps after purification. The fusion protein of SrtA and Tev protease, SrTev, exhibited superior activity compared to separate enzymes, providing a robust approach for efficient protein conjugation.

The exterior modifications involved attaching GFP to the surface of encapsulin nanocompartments without adversely affecting their shape and size, while attaching proteins inside the encapsulin nanoparticles revealed a significant enhancement in the quantity of loaded proteins compared to conventional Cargo-Loading-Peptide (CLP) sequences. However, cargo overload resulted in varied shapes and sizes of encapsulin nanoparticles, deviating from the typical spherical form. This phenomenon suggests the need for careful consideration when aiming for enhanced cargo loading to prevent alterations in nanoparticle morphology. Also, we utilized the developed system to attach an anti-CD3 scFv onto the surface of the encapsulin nanoparticles, successfully targeting Jurkat cells. This serves as a prominent example of the system’s functionality.

Moreover, this study addressed challenges associated with other methods, such as chemical linkers and the SpyTag-SpyCatcher system, emphasizing the advantages of the SrtA-mediated approach. The proposed system demonstrated efficiency in attaching proteins to the encapsulin nanocompartments, presenting opportunities for further exploration in nanotechnology for targeted drug delivery and vaccine development.

While the current research opens promising avenues for the applications and engineering possibilities of encapsulins, future investigations should delve into addressing the observed changes in nanoparticle morphology under cargo overload. Additionally, the potential immunogenic effects of the encapsulins and further optimization of the SrtA-mediated system could contribute to refining the technology for broader biomedical applications.

### Content of the supporting information files

A table for sequences of the proteins, SEC and ion-exchange chromatograms, SDS-PAGE gel image of post-translational modified encapsulins in different temperatures, TEM images of the modified encapsulin nanoparticles.

## Supplementary

**Table S1.**
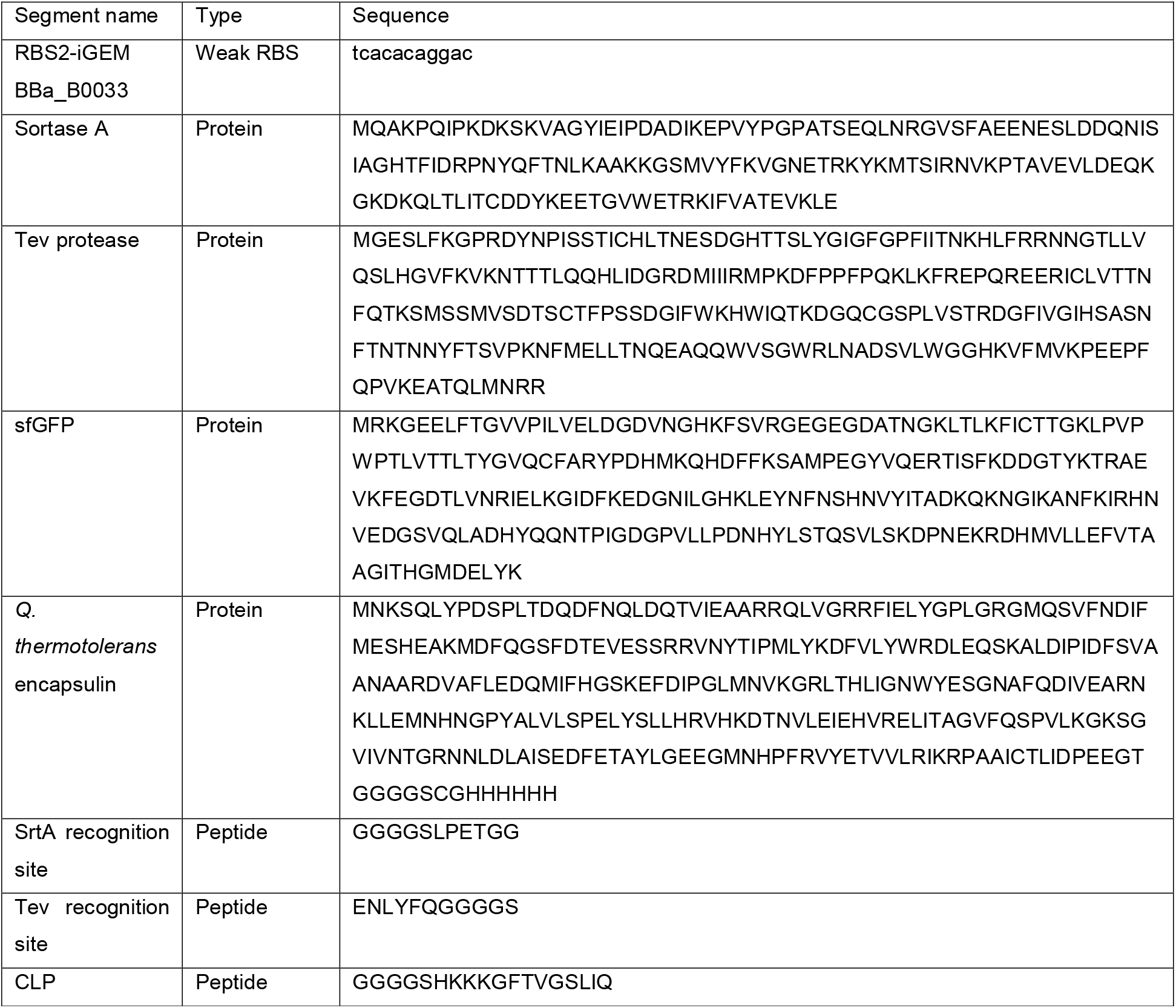
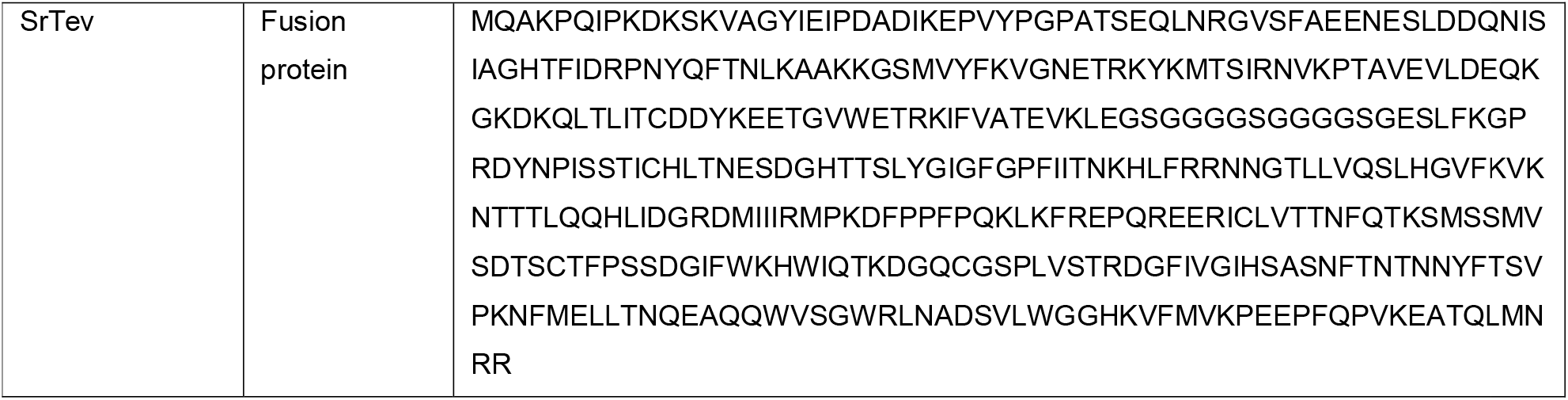

**Figure S1.**
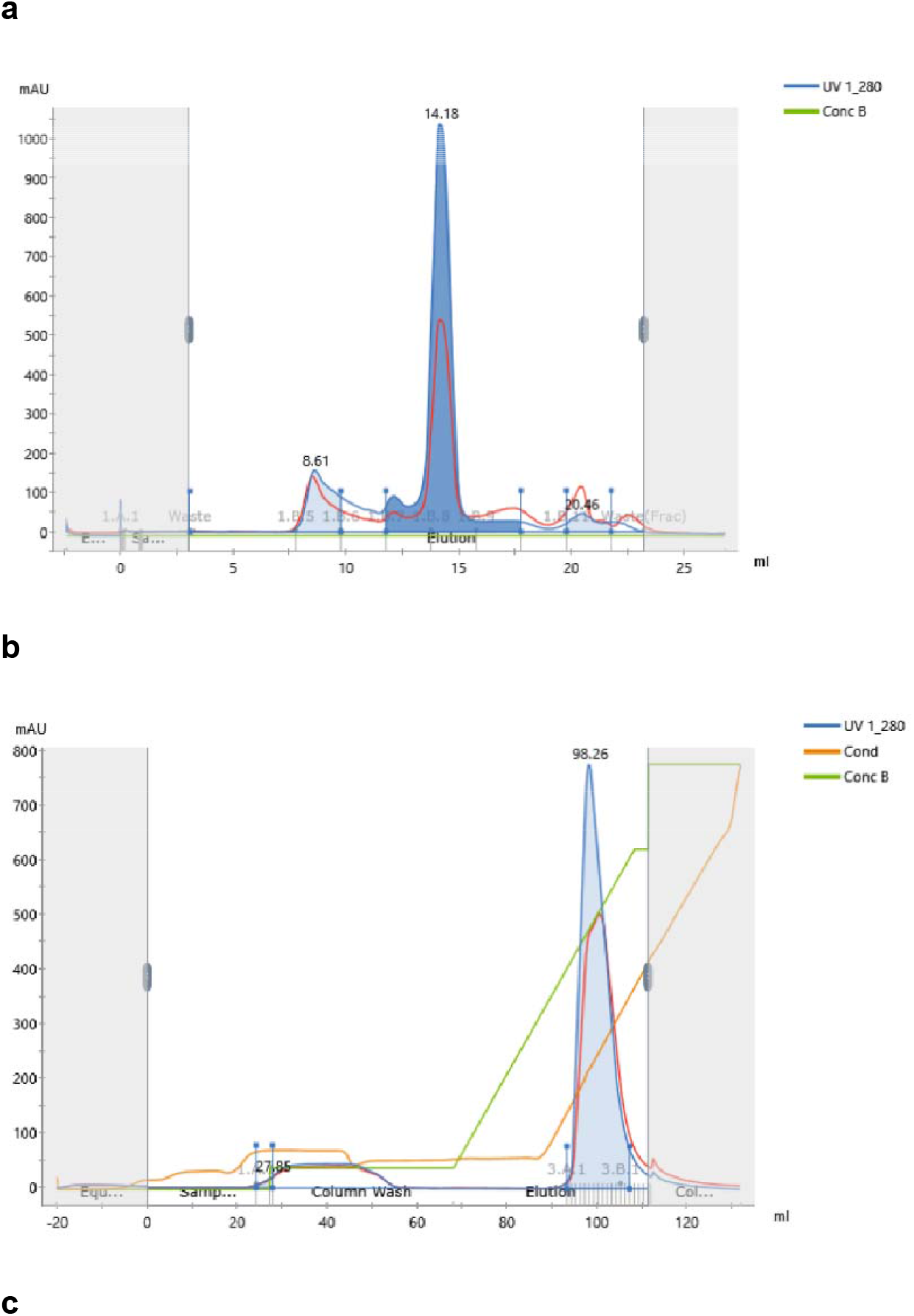
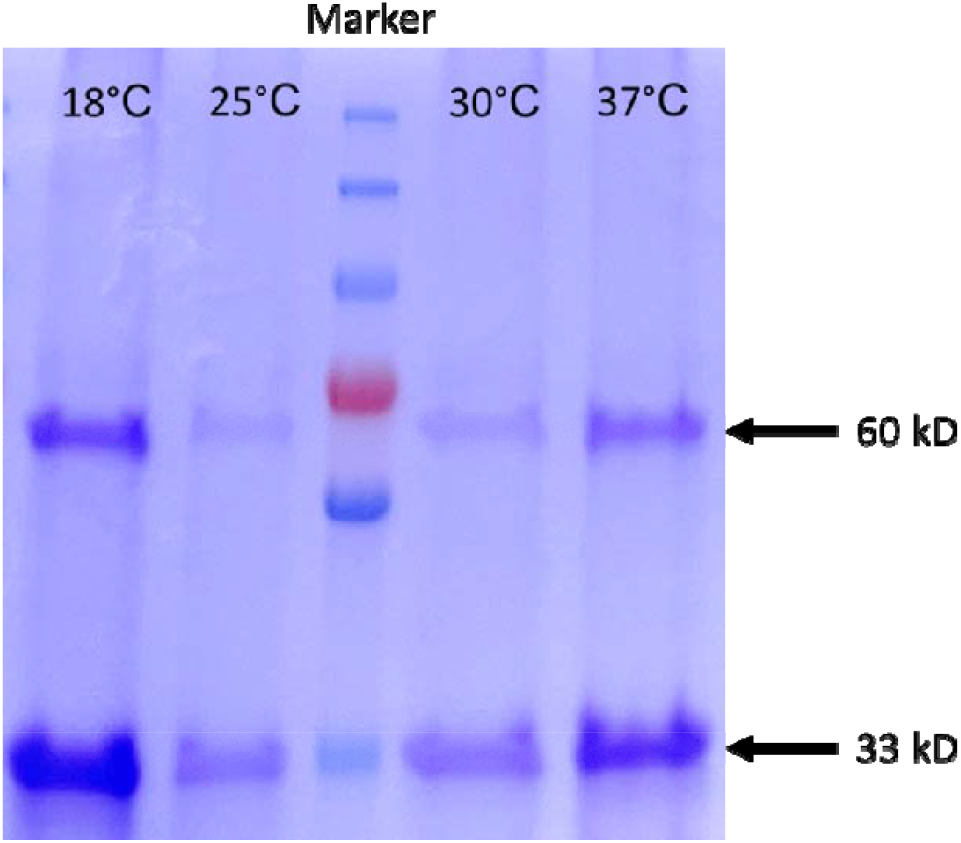
(a) An example of size exclusion chromatogram of encapsulin nanoparticles purification. (b) Ion-exchange chromatogram. (c) SDS-PAGE gel image of expressed encapsulin nanoparticles in different temperatures. This outcome suggests that the sortase-based system effectively facilitates the ligation of the target protein to encapsulin nanoparticles within the temperature range of 18 to 37 degrees Celsius. The 33 kDa bands correspond to encapsulin protomers, while the 60 kDa bands represent ligated protomers.

**Figure S2.**
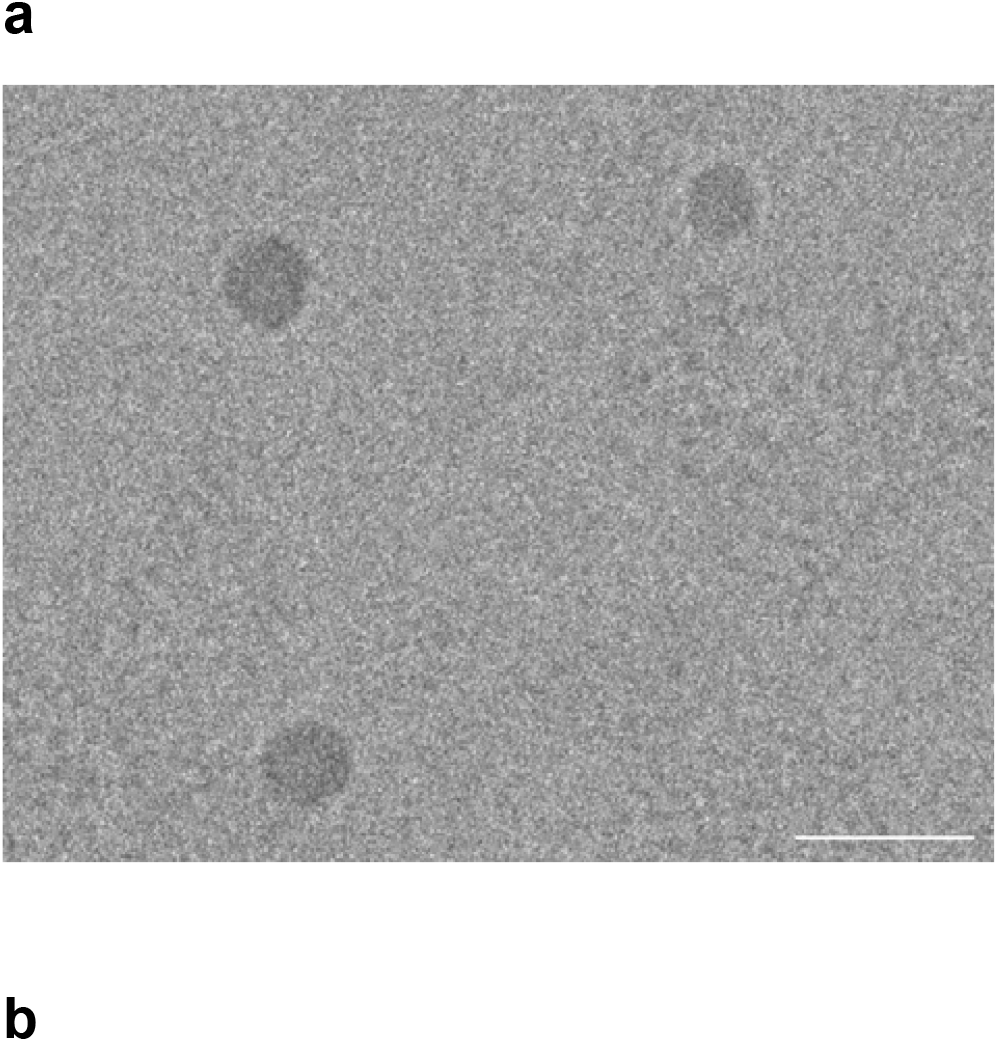
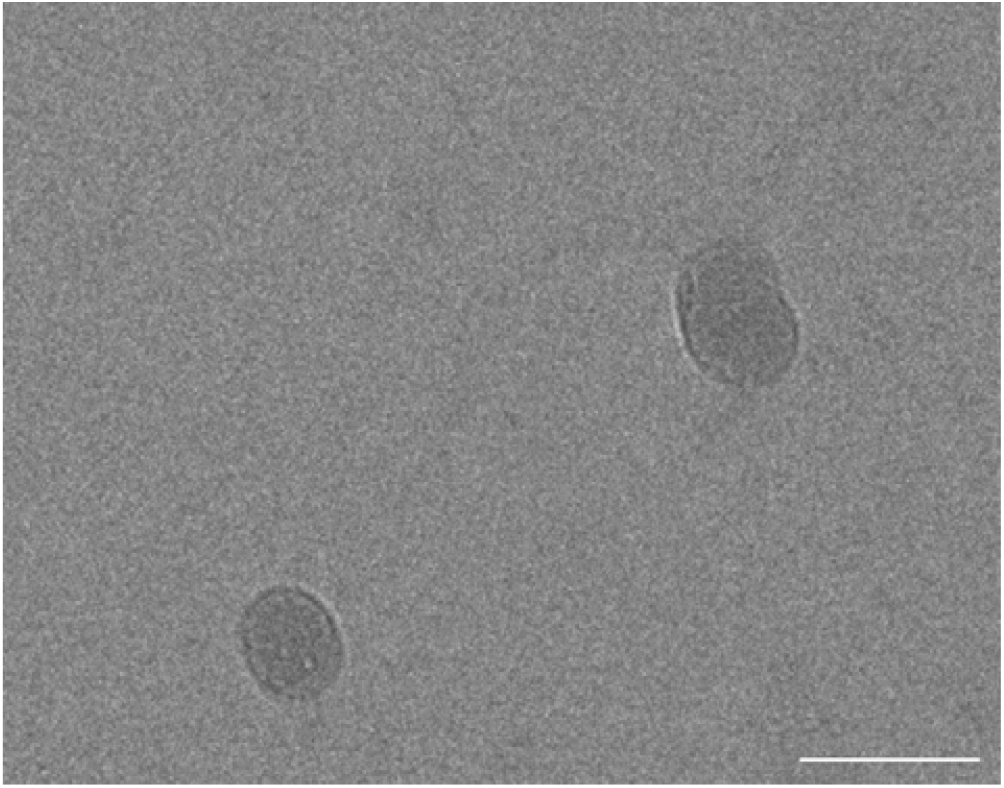
TEM images of the post-translational modified encapsulin nanoparticles. (a) Nanoparticles with GFP proteins attached to their surface displayed a consistent size of 46 nm and uniform shapes. (b) Internal ligation of GFP within encapsulin nanoparticles resulted in diverse shapes and sizes around 76 nm, attributed to cargo overload (GFP). These variations from the typical spherical form are apparent in the image. The scale bar corresponds to 100 nm.

